# Purifying and balancing selection on embryonic semi-lethal haplotypes in a wild mammal

**DOI:** 10.1101/2022.12.02.518882

**Authors:** M.A. Stoffel, S.E. Johnston, J.G. Pilkington, J.M Pemberton

## Abstract

Embryonic lethal mutations are arguably the earliest and most severe manifestation of inbreeding depression, but their impact on wild populations is not well understood. Here, we combined genomic, fitness and life-history data from 5,925 wild Soay sheep sampled over nearly three decades to explore the impact of embryonic lethal mutations and their evolutionary dynamics. We searched for haplotypes which in their homozygous state are unusually rare in the offspring of known carrier parents and found three putatively semi-lethal haplotypes with 27-46% fewer homozygous offspring than expected. Two of these haplotypes are decreasing in frequency, and gene-dropping simulations through the pedigree suggest that this is partially due to purifying selection. In contrast, the frequency of the third semi-lethal haplotype remains relatively stable over time. We show that the haplotype could be maintained by balancing selection because it is also associated with increased postnatal survival and body weight and because its cumulative frequency change is lower than in most drift-only simulations. Our study highlights embryonic mutations as a largely neglected contributor to inbreeding depression and provides a rare example of how harmful genetic variation can be maintained through balancing selection in a wild mammal population.

## Introduction

Most organisms carry a large number of (partially-) recessive deleterious mutations spread throughout their genomes (Charlesworth & Willis, 2009). While their effects are often concealed as heterozygotes, inbreeding increases genome-wide homozygosity and allows harmful alleles to be expressed. This causes a reduction in fitness in the offspring of related parents, a phenomenon termed inbreeding depression (Charlesworth & Willis, 2009). Inbreeding depression in wild populations has mostly been measured on a genome-wide scale, so that little is known about the effect sizes and location of loci involved (Kardos et al., 2016). For small populations, theory predicts that strongly deleterious recessive mutations are rapidly purged because they are often exposed to selection as homozygotes (Hedrick & Garcia-Dorado, 2016). In line with this, recent whole-genome sequencing studies frequently show purging of predicted loss-of-function mutations in small or bottlenecked populations (Xue et al., 2015; Grossen et al., 2020; Khan et al., 2021). However, large effect deleterious mutations sometimes drift to higher frequencies even in small populations due to stochasticity in mating patterns and demography. For example, a single recessive allele causing a lethal form of dwarfism affects the Californian condor (*Gymnogyps californianus*) and segregates at a frequency of 9% (Ralls et al., 2000). Similarly, in Scottish red-billed choughs (*Pyrrhocorax pyrrhocorax*), a recessive mutation causes blindness in 1-6% of nestlings (Trask et al., 2016). Despite their potential importance, strongly deleterious recessive alleles are difficult to detect in wild populations, because they do not usually have an obvious phenotypic effect, are present at very low frequencies, or cause prenatal mortality.

Embryonic lethal mutations that prevent an individual from being born are arguably the earliest and most severe manifestation of inbreeding depression. They are likely to be relatively common, as loss of function mutations are lethal in around one third of mammalian genes, and most of these are probably lethal pre-rather than postnatally (Dickinson et al., 2016; Georges et al., 2019). In farm animals, reverse genetic screens for depleted haplotype homozygosity have identified dozens of embryonic lethals (VanRaden et al., 2011; Fritz et al., 2013; Charlier et al., 2016; Derks et al., 2017; Jenko et al., 2019). These can have substantial effects on the population as a whole, with around 0.5% of embryos being affected by embryonic lethal mutations in cattle and pigs (Charlier et al., 2016; Derks et al., 2019). While different methods exist to detect embryonic lethals and semi-lethals (mortality of some but not all embryos), the most reliable screens identify parents which are known carriers of a focal haplotype and test whether their living offspring are less often homozygous than expected. However, these screens need large sample sizes, dense genomic data, and genetic sampling immediately after birth to exclude postnatal lethality, which has so far largely prevented the detection of embryonic lethal mutations in wild populations.

The Soay sheep of St. Kilda are descendants of early Bronze Age sheep which have roamed the Scottish St. Kilda archipelago freely and unmanaged for thousands of years. For nearly four decades, a part of the population in the Village Bay area of Hirta has been subject to a long-term study with genomic, phenotypic and life-history data collected for thousands of individuals, providing a unique opportunity to shed light on the impact of embryonic lethal mutations in the wild. Here, we scanned high-density SNP genotypes of nearly six thousand Soay sheep for embryonic lethal and semi-lethal haplotypes, explored whether their dynamics over the time are driven by selection or genetic drift and assessed their potential impact on postnatal fitness.

## Results

We searched for haplotypes carrying putatively embryonic-lethal and semi-lethal mutations by screening for depleted haplotype homozygosity in a dataset of 5,925 wild Soay sheep with phased genotypes at 417k autosomal SNPs. Specifically, we identified pairs of parents each carrying at least one copy of a focal haplotype and assessed whether their offspring were less often homozygous for that haplotype than expected. Initially, we tested haplotypes ranging in length from 100 to 500 SNPs (~700Kb to ~3,500Kb). The patterns of homozygous haplotype deficiency were qualitatively similar for different haplotype lengths (Supplementary Figure 1). We therefore subsequently focused on haplotypes with a length of 400 SNPs (~2,800Kb), as all genome-wide significant regions in this analysis were clearly present for all other haplotype lengths (Supplementary Figure 1).

Overall, no putatively fully lethal haplotype reached genome-wide significance, although one haplotype on chromosome 9 (6.64-8.74 Mb) was suggestive, with zero observed homozygotes despite 8.25 expected homozygote offspring from 33 carrier x carrier matings (*χ p-value* = 0.0009, df = 1). We detected three semi-lethal haplotypes (Figure 1), from here on named SEL05 (**S**oay **E**mbryonic semi-**L**ethal Chr. 5; 37.2-39.8 Mb, carrier x carrier matings: N = 800, expected homozygotes: N = 258.50, observed homozygotes: N = 189, *χ^2^ p-value* = 1.49 x 10^-7^, df = 1, SEL07 (Chr.7; 71.2-73.3 Mb, carrier x carrier matings: 382, exp.: 105.75, obs.: 58, *χ*^2^ *p-value* = 1.28 x 10^-8^, df = 1) and SEL18 (Chr.18, 3.23-5.68 Mb, carrier x carrier matings: 815, exp.: 254.25, obs.: 176, *χ^2^ p-value* = 3.29 x 10^-9^, df = 1), with 27%, 47% and 31% fewer homozygous offspring than expected, respectively (Supplementary Table 1). Assuming complete sampling of individuals in the study area, these three semi-lethal haplotypes have therefore potentially prevented around 199 individuals from being born.

**Figure 1:**
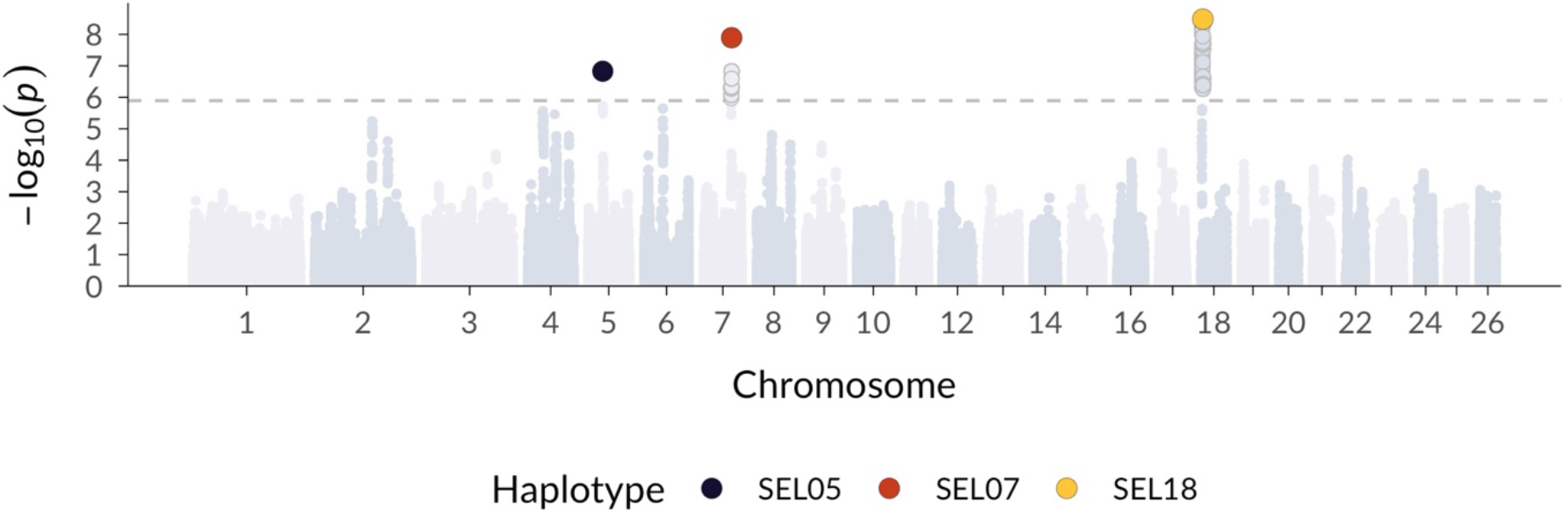
Genome-scan for embryonic lethal haplotypes in Soay sheep. Shown are p-values for a homozygous haplotype deficiency test in the offspring of carrier x carrier matings, in 400-SNP haplotypes sliding one SNP at a time across the genome. The dotted line marks the genome-wide significance threshold.

To better understand the short-term evolutionary dynamics of the semi-lethal haplotypes in the Soay sheep population since 1990, we performed gene-dropping simulations through the pedigree (Figure 2A). This approach allows us to evaluate whether the observed changes in haplotype frequency over time are consistent with expectations from genetic drift alone or whether selection could be contributing factor (MacCluer et al., 1986; Gratten et al., 2012; Johnston et al., 2013). From 1990 to 2018, SEL07 and SEL18 declined in frequency from 19% to 7% and from 32% to 18%, respectively (Figure 2A). The steep decline in frequency of SEL07 is unlikely to have occurred by drift alone, with only 7.4% of simulations resulting in steeper declines (Figure 2A, B). In contrast, there is little evidence for purifying selection in SEL18, as 22.1% of simulations showed steeper frequency declines, indicating that drift alone can frequently result in a decline of this magnitude (Figure 2A, B). Additionally, we explored the potential role of recombination in breaking down the haplotypes at rates that could have led to similar decreases, but found that gene-dropping simulations including recombination yielded qualitatively similar patterns (Supplementary Figure 2).

**Figure 2:**
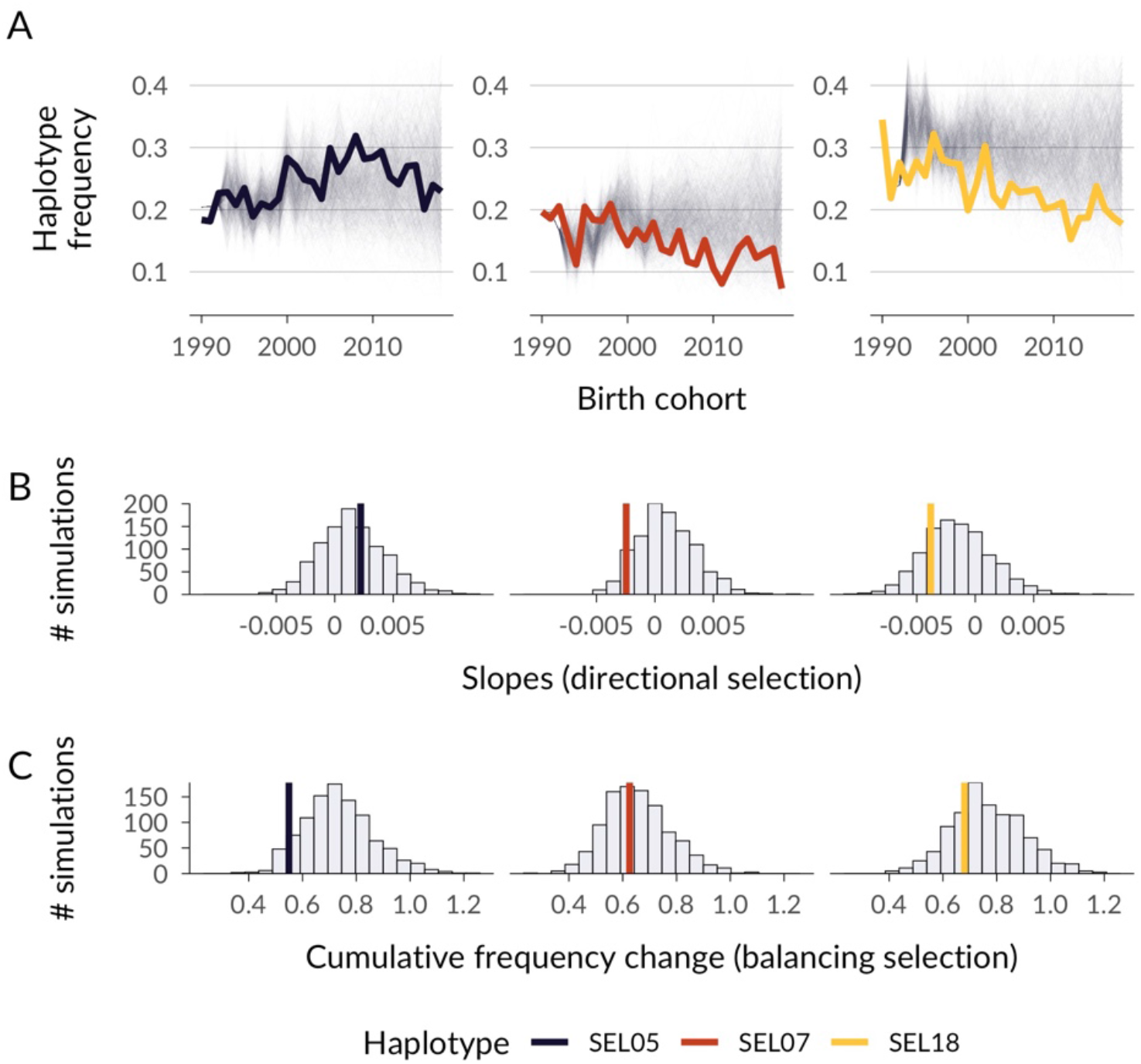
Empirical haplotype dynamics and gene-drop simulations for embryonic semi-lethal haplotypes in Soay sheep. Panel A shows the empirical haplotype frequencies per birth cohort from 1990 to 2018 as thick coloured lines and the results of 1,000 gene-drop simulations through the pedigree as thin grey lines. Gene-drop simulations represent possible frequency changes over time under genetic drift alone. Panel B compares linear model slopes of the empirical haplotype frequencies over time to simulated slopes as an indicator for directional selection. Panel C compares the cumulative frequency change of gene-drop simulations to the empirical haplotype frequency change as an indicator for balancing selection.

In contrast, the frequency of SEL05 did not decline and remained relatively stable over the last decades (from 20% in 1990 to 23% in 2018). This could be due to balancing selection, for example when the semi-lethal mutation is in linkage disequilibrium (LD) with an allele under positive selection.

To test this, we compared the cumulative frequency change seen in gene-drop simulations to the empirical data. Under balancing selection, we would expect the frequency change seen in drift-only gene-drop simulations to be larger than in the empirical data. Only 6.7% of simulations had a lower cumulative frequency change than observed empirically, suggesting that the relative stability in the frequency of SEL05 is unlikely under genetic drift alone (Figure 2C).

Finally, to explore whether embryonic semi-lethal haplotypes impact postnatal fitness, we estimated the effects of having one or two copies of each haplotype on first-year survival using Bayesian generalised linear mixed models. We fitted all three haplotypes simultaneously as predictors and also included other phenotypic and environmental variables in the model (see Methods). Haplotype SEL18 had no effect on first-year survival, while SEL07 showed a tendency to decrease survival in heterozygote individuals, although credible intervals overlapped zero (Figure 4A, Supplementary Table 3), suggesting that deleterious effects of both haplotypes are largely expressed prenatally.

In contrast, SEL05 was associated with an increased first year survival when heterozygous (posterior mean log-odds estimate, 95% credible interval = 0.275, [0.015, 0.539], Supplementary Table 3). This translates into a predicted increase in survival probability of 6.58% (6.58, [0.350, 12.9], Figure 3A) when comparing individuals with one vs. no copy of SEL05 and when holding all other predictors constant at their mean and other haplotypes at their reference levels (0 copies). To examine a potential pathway for how SEL05 could increase survival, we fitted a model of August weight, a key fitness-related trait, with the same predictors as before. In line with higher survival, lambs with one copy of SEL05 were predicted to be 166 grams heavier (posterior mean estimate [95% credible interval] = 0.166 [0.043, 0.289]), and lambs with two copies were predicted to be 212 grams heavier (0.212, [−0.042, 0.466]), although credible intervals were wide due to a relatively small sample size for homozygous individuals (Figure 3B; see Supplementary Table 3 for all model estimates). In contrast there was no association between SEL07 or SEL18 and August weight.

**Figure 3:**
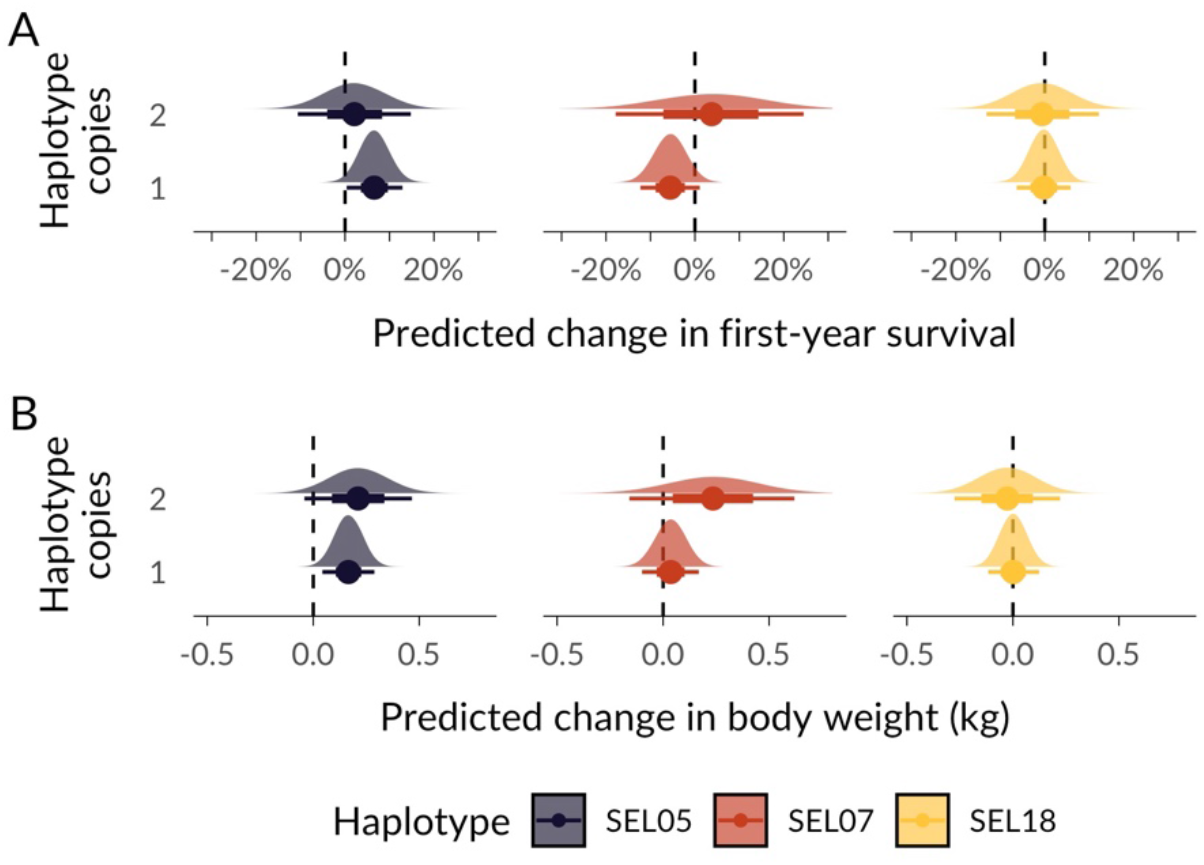
GLMM predicted differences in (A) first-year survival and (B) lamb August body weight for individuals with one and two copies of each haplotype, compared to the reference level of having no copy of the focal haplotype. Fitted models included genotypes for all three haplotypes simultaneously. Half-eye plots show the posterior distribution plus the posterior mean as a point and the 66% and 95% credible intervals as thick and thin lines.

## Discussion

Detecting lethal and semi-lethal mutations in wild populations remains a major challenge, as they are rare and can be lethal even before birth. In this study, we identified three semi-lethal haplotypes linked to mortality in one third up to nearly half of homozygous embryos in a wild population of Soay sheep on the Scottish St. Kilda archipelago. Notably, homozygous haplotype carriers in the (living) population did not suffer from reduced survival, suggesting that the harmful effects are specific to embryo development. Over the last two decades, purifying selection is likely to have contributed to a reduction in the frequency of at least one these haplotypes (SEL07) in the population. In contrast, the third semi-lethal haplotype (SEL05) is relatively stable over the recent past. Gene-drop simulations and an association with increased survival and body weight in lambs suggest that the haplotype frequency is partially maintained by balancing selection.

All three embryonic semi-lethal haplotypes were present at relatively high frequencies between 19% and 32% in the birth cohort of 1990. This is not surprising, as genetic drift is strong in the Soay population. The estimated N_e_ is only around 200 individuals (Kijas et al., 2012), and the population experienced a recent bottleneck, where 85 sheep including 20 males were transferred from the island of Soay to the island of Hirta in 1934-5, founding the population which we now study (Clutton-Brock & Pemberton, 2004). Therefore, the founder event and demographic stochasticity after the bottleneck could have led to a rise in the frequency of strongly deleterious mutations. A third explanation for semi-lethal mutations at high frequencies is a possible admixture event around 150 years ago with the now extinct Dunface breed, which could have introduced deleterious variation into the population (Feulner et al., 2013). Finally, while the three detected haplotypes had relatively high frequencies, we expect this to be an ascertainment bias due to limited statistical power, where most semi-lethals and lethals remain undetected as they were simply too rare to reach genome-wide significance in our haplotype scan. Consequently, while strongly deleterious mutations are generally expected to be purged when N_e_ is small (Hedrick & Garcia-Dorado, 2016), their potential impact should not be ignored in real world populations, where demographic stochasticity and genetic drift can be high.

Over the last 25 years, the frequencies of the semi-lethal haplotypes SEL07 and SEL18 declined in the population, as would be expected if purifying selection is effective. However, in small populations, genetic drift can substantially change allele frequencies even in the absence of selection. Using gene-drop simulations based on the Soay sheep pedigree, we established a baseline expectation for haplotype frequency changes under drift alone. Only 7.4% of simulations showed steeper declines for SEL07 than observed empirically, suggesting that purifying selection may be contributing to the decline. Moreover, SEL07’s frequency decreased by 12% in less than ten generations, which suggests that selection can be effective in reducing strongly deleterious variation within short, ecological timescales. The efficient selection against SEL07 in the Soay population is consistent with theoretical (Hedrick & Garcia-Dorado, 2016) and empirical (Grossen et al., 2020; Khan et al., 2021; Stoffel et al., 2021a) research showing that inbreeding depression in small populations is more likely to be a consequence of many weakly rather than fewer strongly deleterious alleles.

Surprisingly, haplotype SEL05 had a relatively stable population frequency of around 20% over the last two decades despite its putative embryonic semi-lethality, and further analyses showed some support for balancing selection. Comparing SEL05 to drift-only gene-drop simulations, we showed that 93% of simulations had a higher cumulative frequency change than SEL05, making SEL05 more stable than expected in drift-only scenarios. Moreover, SEL05 was positively associated with postnatal fitness. Lambs which were heterozygous (but not homozygous) for SEL05 had a 6% higher predicted survival probability over their first winter. A second analysis of August body weight provided a potential pathway, as lambs with one or two copies of the haplotype were 166 and 212 grams heavier when controlling for other predictors such as skeletal size (hindleg length) and inbreeding coefficient. There are several mechanistic explanations why SEL05 could be under balancing selection. One is antagonistic pleiotropy, where the same genetic variant has opposing effects on fitness, and which has been suggested as a widespread mechanism maintaining deleterious alleles (Carter & Nguyen, 2011). In farm animals for example, embryonic lethal mutations are maintained at high frequencies due to pleiotropic effects on milk yield in cows and growth in pigs (Kadri et al., 2014; Derks et al., 2018). Another explanation is linkage disequilibrium (LD) between the semi-lethal mutation and an allele under positive selection that increases body weight and survival. LD stretches over long distances in Soay sheep, with a half-decay around 600Kb (Stoffel et al., 2021a), and analysing relatively long haplotypes makes it more likely to pick up antagonistic alleles too. To sum up, haplotype SEL05 was associated with both prenatal semi-lethality and higher postnatal weight and survival. Its frequency was also unusually stable over the last decades, all of which suggests that it is maintained by balancing selection.

Lastly, our study raises the question of how much embryonic lethal and semi-lethal alleles collectively contribute to inbreeding depression in natural populations. If homozygous carriers are absent or rare in the living population, the effects of embryonic lethal alleles will be largely neglected in estimates of inbreeding depression based on postnatal fitness. While some animals might be able buffer the fitness effects of lost embryos through re-mating, there could be a substantial population-wide impact, especially in small populations where carrier frequencies of specific mutations can be high. Currently, genome-wide scans for depleted homozygosity are not feasible in most wild populations due to the need for large sample sizes, extensive parentage information and dense genomic data. A promising avenue is a two-step approach, in which genome-sequence based predictions of loss-of-function mutations could limit the number of target regions, and thereby increase the power to detect depleted homozygosity and embryonic lethals. Overall, our study reveals the potential contribution of semi-lethal mutations to inbreeding depression and individual fitness and highlights balancing selection as a mechanism for the maintenance of harmful genetic variation in wild populations.

## Materials and Methods

### Study population

Soay sheep are descendants of primitive European domestic sheep and have lived unmanaged on the St. Kilda archipelago, Scotland, for thousands of years (Clutton-Brock & Pemberton, 2004). A part of the population in the Village Bay area on the island of Hirta (57 49’N, 8 34’W) has been the focus of a long-term individual-based study since 1985 (Clutton-Brock & Pemberton, 2004). More than 95% of individuals in the study area are ear-tagged within a week after birth during the lambing season from March to May, and DNA was extracted from either blood samples or ear punches. In order to impute genotypes, we assembled a pedigree based on 431 unlinked SNP markers from the Ovine SNP50 BeadChip using the R package Sequoia (Huisman, 2017). In the few cases where no SNP genotypes were available, we assigned parents either from field observations or microsatellite markers (Morrissey et al., 2012). All animal work was carried out according to UK Home Office procedures and was licensed under the UK Animals (Scientific Procedures) Act of 1986 (Project License no. PP4825594).

### Fitness and phenotype data

Routine mortality checks, in particular during peak mortality in February, usually find around 80% of deceased animals (Bérénos et al., 2016). Here, we analysed 1) ‘first year survival’, where every individual was given a 1 if it survived from birth (March to May) to the 30^th^ April of the next year, and a 0 if it did not, with measures available for 5,925 individuals born from 1 979 to 2018. We also used phenotypic measures for lamb body weight in kg (to the nearest 0.1kg) and lamb hindleg size in mm (to the nearest mm), both of which are measured in lambs every August.

### Genotyping

We genotyped a total of 7,700 Soay sheep on the Illumina Ovine SNP50 BeadChip resulting in 39,368 polymorphic SNPs after filtering for SNPs with minor allele frequency > 0.001, SNP locus genotyping success > 0.99 and individual genotyping success > 0.95. We then used the *check.marker* function in GenABEL version 1.8-0 (Aulchenko et al., 2007) with the same thresholds, including identity by state with another individual < 0.9. We also genotyped 189 sheep on the Ovine Infinium HD SNP BeadChip, resulting in 430,702 polymorphic SNPs for 188 individuals, after removing monomorphic SNPs, and filtering for SNPs with SNP locus genotyping success > 0.99 and individual sheep with genotyping success > 0.95. These sheep were specifically selected to maximise the genetic diversity represented in the full population (for full details, see Johnston, Bérénos, Slate, & Pemberton, 2016). All SNP positions were based on the Oar_v3.1 sheep genome assembly (GenBank assembly ID GCA_000298735.1 (Jiang et al., 2014)).

### Genotype imputation and phasing

The detailed genotype imputation methods are presented elsewhere (Stoffel et al., 2021a). Briefly, we first merged the datasets from the 50K SNP chip and from the HD SNP chip with the function --bmerge in PLINK v1.90b6.12 (Purcell et al., 2007), resulting in a dataset with 436,117 SNPs including 33,068 SNPs genotyped on both SNP chips. We then discarded SNPs on the X chromosome and focused on the 419,281 SNPs located on autosomes. To impute SNPs with genotypes missing in individuals genotyped at the lower SNP density, we used AlphaImpute v1.98 (Hickey et al., 2012), which uses both genomic and pedigree information for phasing and subsequent imputation of missing genotypes. After imputation, we filtered SNPs with call rates below 95%. Overall, this resulted in a dataset with 7691 individuals, 417,373 SNPs and a mean genotyping rate per individual of 99.5% (range 94.8%-100%). We evaluated the accuracy of genotype imputation using 10-fold leave-one-out cross-validation. In each iteration, we randomly chose one individual genotyped on the high-density (HD) SNP chip, masked genotypes unique to the HD chip and imputed the masked genotypes. This allowed us to compare the imputed genotypes to the true genotypes and to evaluate the accuracy of the imputation. Overall, 99.3% of genotypes were imputed correctly. To conduct haplotype-based analyses, we phased the imputed SNP dataset using SHAPEIT4 (Delaneau et al., 2019) using the Soay sheep linkage map (Johnston et al., 2016) and default parameter values. To infer linkage map positions for imputed SNPs, we used interpolation by assuming a constant recombination rate in genomic regions between linkage mapped SNPs (Stoffel et al., 2021b).

### Homozygous haplotype deficiency analyses

We identified haplotypes with putatively recessive (semi-)lethal mutations by testing whether offspring of carrier x carrier matings were less often homozygous for a given haplotype than expected. Specifically, for a focal haplotype *h*, we first identified parent-offspring trios where both parents carried at least one copy of *h*. We then calculated the expected number of homozygous offspring with 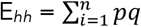 where *n* is the number of parent pairs, *p* is the transmission probability of haplotype *h* for the female and *q* is the transmission probability of haplotype *h* for the male. Transmission probabilities are 0.5 if the individual is heterozygous and 1 if it is homozygous for *h*. Based on the observed number of homozygous individuals O*_hh_* we then followed Jenko *et al*. (2019) and calculated a one way and one degree of freedom Chi square test statistic *χ*^2^ = (Ohh*_hh_* - E*_hh_*)^2^ / E*_hh_* + (O*_non-hh_* - E*_non-hh_*)^2^ / E*_non-hh_* with *non-hh* being the number of offspring which is either heterozygous or contains two copies of alternative haplotypes. To scan the genome for haplotypes deficient in homozygotes we used overlapping windows with varying length (100-500 SNPs) sliding one SNP at a time across the autosomal genome. For example, for the haplotype length of 100 SNPs we started with a window ranging from SNP 1 on chromosome 1 to SNP 100 on chromosome 1, identified all existing haplotypes with frequencies above 0.1 % in the population in this window, and then conducted the test for each identified haplotype. In line with previous work on high-density SNPs in Soay sheep (Stoffel et al., 2021a) we used a genome-wide significance threshold of *p* < 1.28 * 10^-6^, which is a Bonferroni corrected p-value based on the number of independent tests (n_eff_ = 39,184) estimated using SimpleM (Gao et al., 2008) which takes into account linkage disequilibrium between markers. The threshold is not statistically precise, because it is difficult to determine the exact independent number of tests for a haplotype-based sliding window analysis. Per genomic window there are usually more than two haplotypes, so we evaluate more tests per region compared to a biallelic SNP-based association study. However, haplotypes are not independent as they overlap substantially when sliding them over the genome SNP by SNP. The genome-wide significance threshold should therefore be interpreted cautiously. Finally, to explore the effects of haplotype length on detecting homozygosity deficiency, we re-ran the genome scan with haplotype lengths ranging from 100 to 500 SNPs.

### Gene-drop analysis

We tested whether haplotype frequency changes across time are in line with genetic drift in the Soay sheep pedigree or potentially the result of selection using gene-drop simulations in genedroppeR v0.1.0 (code available at https://github.com/susjoh/genedroppeR).

Each individual present in the Soay sheep pedigree was assigned to a cohort based on their birth year. All cohorts from 1990 onwards were included, as the proportion of individuals genotyped before this time was below 70%. Then, the proportion of individuals defined as “founders” in each cohort (i.e. both parents are unknown) were determined; visual observation indicated that proportion of founder individuals declined rapidly from 1990 to 1992; these three cohorts are hereafter defined as the “sampled” cohorts, with the cohorts from 1993 to 2018 defined as the “simulated” cohorts. A total of 1,000 gene-drop simulations were conducted as follows. For all founder individuals in the sampled cohorts, haplotypes were sampled with the probability of their observed frequency in the individual’s cohort. For non-founder individuals in the sampled cohorts, a haplotype was sampled from each of its parents assuming Mendelian segregation (Pr = 0.5); if one parent was missing, then a haplotype was sampled as for the founder individuals above. In the simulated cohorts, non-founder individuals sampled a haplotype from each parent assuming Mendelian segregation (Pr = 0.5). The haplotype frequencies were then calculated within each cohort. Finally, for any founder individuals or those with a missing parent in the simulated cohorts, haplotype(s) were sampled based on the haplotype frequencies in the rest of the simulated cohort. This generated simulated genotypes for each individual in the pedigree, which could then be used to generate a null distribution of haplotype frequencies and their changes over time (i.e. as expected under genetic drift alone) across the simulated cohorts (from 1993 to 2018). Comparisons were made only using individuals with known genotypes to allow a direct comparison between observed and simulated data. Using these data, we examined two aspects of allele frequency change over time using cohort year as a linear variable:

1. Directional selection: For each simulation, we modelled the frequency change the focal haplotype over time using a linear regression. The probability of observing the true slope under drift was determined by comparing it to the distribution of simulated slopes from 1993 to 2018;
2. Balancing selection: For each simulation, we modelled the cumulative change of the focal haplotype (i.e. the sum of the differences between allele frequencies from year to year) using a linear regression. Here, we assume that alleles with lower cumulative change from 1993 to 2018 may be subject to balancing selection.

### Modelling

We estimated the effects of semi-lethal haplotypes on postnatal traits using Bayesian generalised linear mixed models (GLMM) in brms v2.15.0 (Bürkner, 2017), a high-level R interface to Stan (Carpenter et al., 2017). For all models, we used a normal prior with mean = 0 and standard deviation = 5 for population-level (fixed) effects and the default half Student-*t* prior for the standard deviation of group-level (random effects) parameters. We ran four MCMC chains with the NUTS sampler with 10,000 iterations each, a warmup of 5,000 iterations and no thinning. All chains were visually checked for convergence and the Gelman-Rubin criterion was < 1.1 for all predictors, indicating good convergence (Gelman & Rubin, 1992).

### Survival analysis

In the first model, we estimated the effects of semi-lethal haplotypes on first-year survival using a binomial model with logit link. We fitted first-year survival as a response variable and genotype dosages for the three haplotypes as predictors, with values 0 = two copies of alternative haplotypes, 1 = one copy of the focal haplotype and 2 = homozygous for the focal haplotype. These genotypes were fitted as factors, so that the model estimates differences between the reference level (2 alternative haplotypes) and 1 or 2 copies of the focal haplotype, respectively. Specifically, we used the following model structure based on n = 2294 complete observations: using the following model:

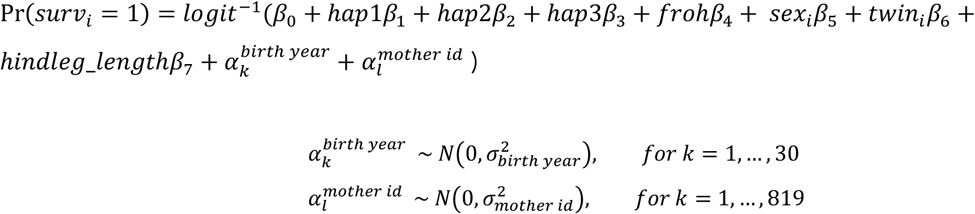

The probability of survival for observation *i* (Pr(*surv_i_* = 1)) was modelled with an intercept *β*_0_, seven population level (fixed) effects, which estimate the effects of the three haplotypes, individual inbreeding coefficient F_ROH_ calculated as the sum of runs of homozygosity (ROH) > 1Mb divided by the autosomal genome size (see Stoffel et al., 2021a for details), sex of the individual (female = 0, male = 1), whether it was a twin (no = 0, yes = 1), and an individual’s skeletal size via its August hindleg length. The latter was fitted to control for variation in individuals due to when they are born in a given year, as smaller individuals that are born later have a smaller chance of surviving the winter. The model also included two group-level (random) intercept effects for birth year and maternal identity to model environmental variation across years and maternal effects, respectively. Both F_ROH_ and hindleg length were standardized (z-transformed).

### Body weight analysis

We estimated the effects of semi-lethal haplotypes on body weight (in kg) in lambs using a model with Gaussian error distribution. We fitted the model with the same fixed and random effects and transformations as above, with n = 2286 complete observations:

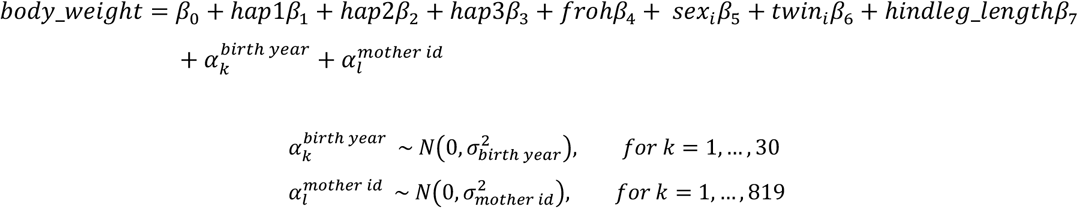

## Acknowledgements

We thank the National Trust for Scotland for permission to work on St. Kilda and QinetiQ, Eurest and Kilda Cruises for logistics and support. We thank Ian Stevenson and many volunteers who have helped with data collection and management and all those who have contributed to keeping the project going. SNP genotyping was conducted at the Wellcome Trust Clinical Research Facility Genetic Core. This work has made extensive use of the Edinburgh Compute and Data Facility (http://www.ecdf.ed.ac.uk/). We are grateful for discussions with with the Wild Evolution Group at the University of Edinburgh, Joel Pick, and especially Janez Jenko. The project was funded through an outgoing Postdoc fellowship from the German Science Foundation (DFG) awarded to MAS and a Leverhulme Grant (RPG-2019-072) awarded to JMP and SEJ. Field data collection has been supported by NERC over many years, and most of the SNP genotyping was supported by an ERC Advanced Grant to JMP.

## Author contributions

JMP and MAS designed the study. JGP is the main Soay sheep project fieldworker and collected samples and life history data. JMP has run the Soay sheep long-term study and organised the SNP genotyping. SEJ wrote the genedroppeR package and built the fundamental genomic database, including genotyping, quality control and linkage mapping. MAS conducted data analyses and drafted the manuscript. MAS, JEP and SEJ jointly contributed to concepts, ideas and revisions of the manuscript.

## Data and code accessibility

All data underlying the analyses are publicly available on Zenodo (Stoffel et al., 2021c). The analysis scripts are available on GitHub (https://github.com/mastoffel/haplotype_homozygosity).

## Notes

### Competing Interest Statement

The authors have declared no competing interest.

