## Supplementary Material for "Purifying and balancing selection on embryonic semi-lethal haplotypes in a wild mammal"

Authors names and addresses:

Stoffel, M.A.[1*], Johnston, S.E.[1], Pilkington, J.G.[1], Pemberton, J.M[1]

[1]Institute of Evolutionary Biology, School of Biological Sciences, University of Edinburgh, Edinburgh, EH9 3FL, United Kingdom

Short running title:
Embryonic semi-lethal mutations in a wild mammal

* Corresponding author:
Martin A. Stoffel
Postal address: Institute of Ecology and Evolution, University of Edinburgh, Edinburgh, EH9 3FL, UK

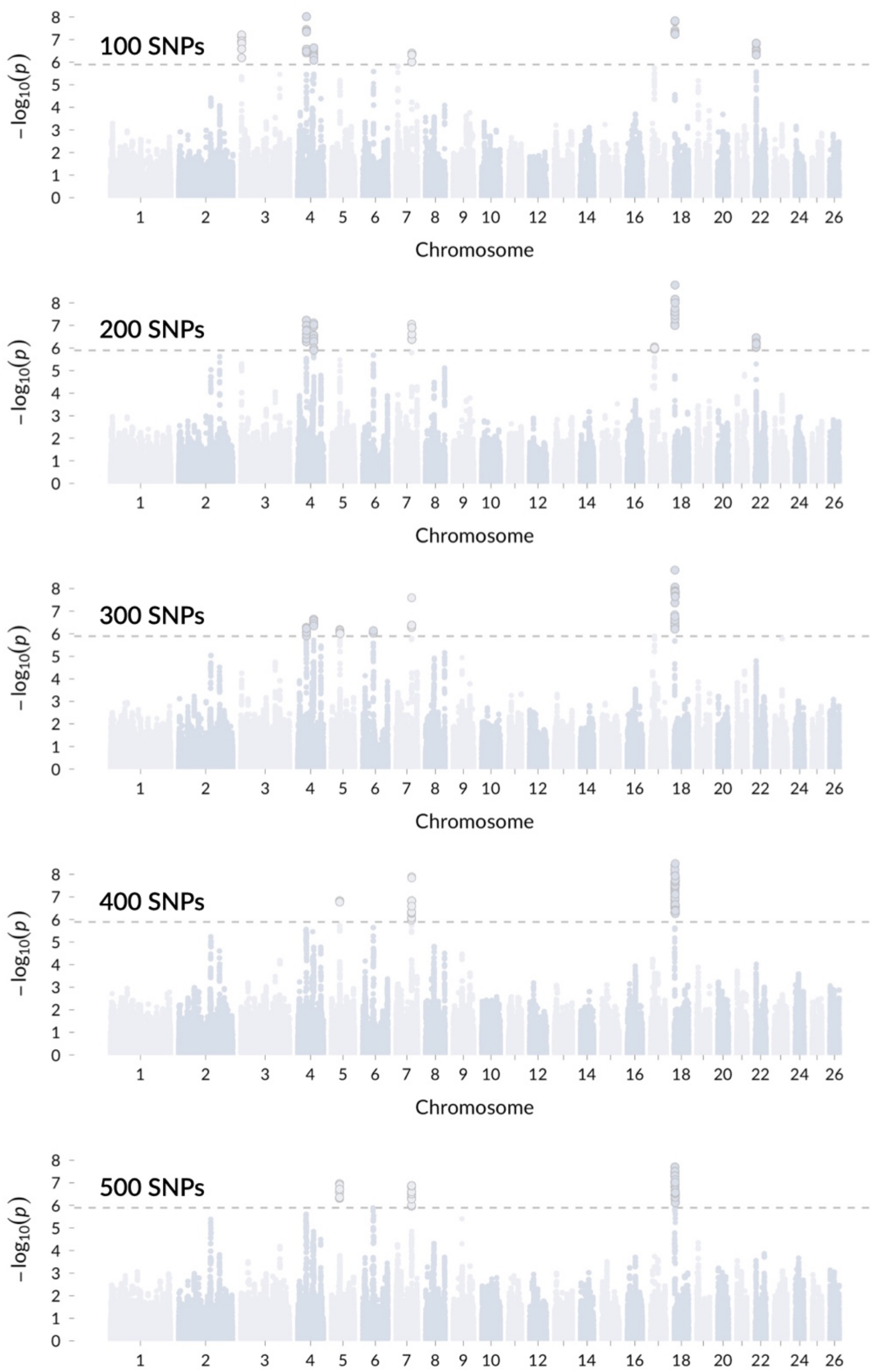


Supplementary Figure 1: Genome-wide scans for depleted haplotype homozygosity in offspring of carrier x carrier matings for different haplotype lengths (100-500SNPs).

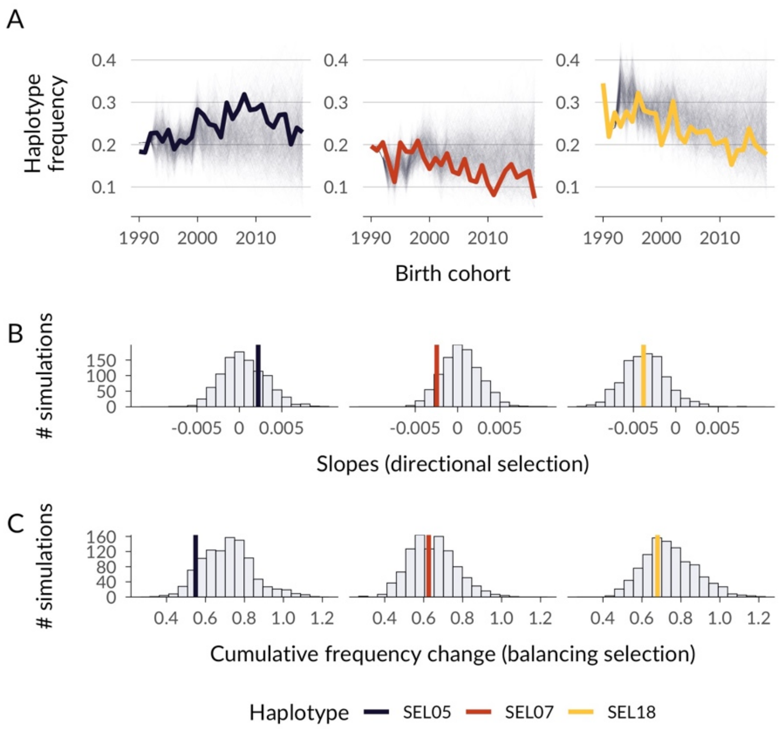


Supplementary Figure 2: Results of gene-dropping simulations with recombination. The genetic map length for each haplotype was determined using the interpolated high-density genetic map from Stoffel, Johnston, Pilkington, & Pemberton, (2021), resulting in map lengths of 2.1, 1.9, and 4.4 cM for SEL05, SEL07, and SEL18, respectively. Cross-overs were simulated on these haplotypes in 2.1%, 1.9%, and 4.4% of meioses, respectively. Panel B shows that this resulted in 74%, 10%, and 47% of slopes being steeper than observed empirically for SEL05, SEL07, and SEL18. Panel C shows that it also resulted in 10%, 46%, and 34% of simulations being more stable than SEL05, SEL07, and SEL18. For simplicity, we did not simulate situations where a focal haplotype re-emerges due to recombination, which is not unlikely given the relatively limited genetic diversity in the Soay population.

| Haplotype | p value (chi-square test) | # observed homozyous offspring | # expected homozygous offspring | # carrier x carrier matings | chromosome | first SNP / last SNP | start position / end position (bp) |
|---|---|---|---|---|---|---|---|
| AAGGACAGGGACAAAGGAGAGGAAAGAGGACAGAGAGAAAAGAGACGGAG AAGGAGGGGCAGGAGCGAAACGGAACAGAAGGGAGAAGAAAAAGGAAAGA GAAGCAAGAAGAGAGGAAAAGAGAGAGAGAGCAAAGAGGAGGAGGAACGA GAGAAAAGAAAAACGAGACGGAGCACAGGGGGGAGAAGGCACGAAGGGAG GGAGCCACGGGAAAAGAGAAGGGAAGAAAAGGAGCGGAAGAGGGGGAAGC GAGCAGAGAGGAGGGAGAAGGGACGAGGAAAACAAGAACGAAGAGGAAGA GAGGAGCAGGGGAGACAAAAGGGAGAACGAAACGAGAAGGGAAAGAGACG CGGAAAAAAGAAAACGAGAAGAGGAGAAGGAAAGAAAAAAAAAAAAAAAGG | 3.29e-09 | 176 | 254.25 | 815 | 18 | oar3_OAR18_4691643-oar3_OAR18_7448353 | 4691643-7448353 |
| GACGAACAAAGGGAAAGAAGAAGAAAAAGAAAAGAGAGGAAGAGCGAGAG GAGGAGAAAAGAGACAAGAGAGGAAAGGAGGGAGCGAAGGGAGAAAAAGA GAGAGGAGGCAAAAAAGACGAGAAAGGAAAGGCGAGCGCAAAAAGGAGAA GAGGAAGAGAGAGAGAGCAAAGAAAGAGAACCAGAGGAAGAGAGAAGACA AGGAAGAAAGAAAAACAAAGGAAAAAAGAGAGGGCGAGGAGGGGGAGAAA AGAAGAAAAAGGGAGACGAAAAGGAACAGGGAGAAACAAGAAGACAAAAA CAAGAAGCGGAATGGAAAGAACGAAGCCGAAAACGAGAGACGAAGAGAAA GGGAAAGGAGAAAACAAACAACACCCGAGCGAGAGAAAAGAGAACGAAGG | 1.28e-08 | 56 | 105.75 | 382 | 7 | oar3_OAR7_71164579-oar3_OAR7_73326795 | 71164579-73326795 |
| AGCGGACAAAGGAGAAAGAGGAAAAAACAGAAAACGAAAGAGCGCAGAAC GCAGAAAGAGGCCGGGCGGCGAAAAACGAGGGGGGGCGGGACGAAAGAAG GAGGAAGGGCAAGCAGAGGCAAAAGAAGAGTGAGGAAAAAAGGAAAAGAA AAGAGGACAAGAAAAAAAAGGAGAGGAAGGAGAAACAAAGAGGAGGGGAA AGGGGAGGGAAGGAAAACTAGCAAAGAACAAAGAAGAGGAGGGAAAGGAA AGAGGAGCACAGGGGGAGAGGAGGGAGAAGAAACGGGAAAGGGACAGAAA GCGCACGAAAGCGGAGGGGAAAGAAAGGGACAAAGAAGGGGAGGGAGCAA GGGAGAAGGAGAGGGAGAAAGAGGGAAGAAAAAGGGAGCGAAGAGAAAAA | 1.49e-07 | 189 | 258.50 | 800 | 5 | oar3_OAR5_37164925-OAR3_212455420.1 | 37164925-39808507 |

Supplementary Table 1: Putatively semi-lethal haplotypes. Shown are the top hits from a genome-scan for depleted haplotype homozygosity in Soay sheep, their chi square p-values, the number of observed and expected homozygous offspring and the overall number of mating pairs as well as the location of the haplotypes in the sheep Oar_v3.1 sheep genome assembly. Note that 'Haplotype' shows the alleles at consecutive SNPs on the SNP chip, and not the genome-sequence.

| Term | Post.Mean | CI (2.5%) | CI (97.5%) | Info |
|---|---|---|---|---|
| Intercept | 0.46 (1.584) | -0.367 (0.693) | 1.31 (3.705) | |
| Population level/fixed effects | | | | |
| SEL05 (1 copy) | 0.275 (1.316) | 0.015 (1.015) | 0.539 (1.715) | categorical |
| SEL05 (2 copies) | 0.089 (1.093) | -0.453 (0.636) | 0.619 (1.857) | categorical |
| SEL07 (1 copy) | -0.236 (0.79) | -0.521 (0.594) | 0.045 (1.046) | categorical |
| SEL07 (2 copies) | 0.159 (1.172) | -0.783 (0.457) | 1.106 (3.021) | categorical |
| SEL18 (1 copy) | -0.009 (0.991) | -0.265 (0.767) | 0.243 (1.275) | categorical |
| SEL18 (2 copies) | -0.026 (0.974) | -0.557 (0.573) | 0.51 (1.665) | categorical |
| $F_{ROH}$ | -0.08 (0.923) | -0.205 (0.814) | 0.044 (1.045) | z-transformed (x-mean(x))/sd(x) |
| Hindleg length | 0.636 (1.89) | 0.486 (1.625) | 0.793 (2.21) | z-transformed (x-mean(x))/sd(x) |
| Sex | -0.966 (0.381) | -1.231 (0.292) | -0.703 (0.495) | categorical (0=female, 1=male) |
| Twin | -0.569 (0.566) | -0.921 (0.398) | -0.215 (0.806) | categorical (0=singleton, 1=twin) |
| Group level/random effects (standard deviation) | | | | |
| Birth Year | 2.217 (9.183) | 1.666 (5.289) | 2.957 (19.246) | n = 30 |
| Mother ID | 0.766 (2.15) | 0.493 (1.636) | 1.025 (2.787) | n = 819 |

Supplementary Table 2: Bayesian GLMM estimates for first-year survival. The sample size was n = 2294. The table shows the posterior mean and 95% credible intervals and information on how each variable was encoded and transformed prior to modelling.

| Term | Post.Mean | CI (2.5%) | CI (97.5%) | Info |
|---|---|---|---|---|
| Intercept | 12.843 | 12.629 | 13.058 | |
| Residual | 1.264 | 1.22 | 1.309 | |
| Population level/fixed effects | | | | |
| SEL05 (1 copy) | 0.166 | 0.043 | 0.289 | categorical |
| SEL05 (2 copies) | 0.212 | -0.042 | 0.466 | categorical |
| SEL07 (1 copy) | 0.035 | -0.1 | 0.169 | categorical |
| SEL07 (2 copies) | 0.237 | -0.159 | 0.631 | categorical |
| SEL18 (1 copy) | 0.002 | -0.117 | 0.123 | categorical |
| SEL18 (2 copies) | -0.027 | -0.276 | 0.223 | categorical |
| $F_{ROH}$ | -0.048 | -0.108 | 0.011 | z-transformed (x-mean(x))/sd(x) |
| Hindleg length | 2.234 | 2.167 | 2.301 | z-transformed (x-mean(x))/sd(x) |
| Sex | 0.608 | 0.492 | 0.726 | categorical (0=female, 1=male) |
| Twin | -0.446 | -0.611 | -0.276 | categorical (0=singleton, 1=twin) |
| Group level/random effects (standard deviation) | | | | |
| Birth Year | 0.477 | 0.346 | 0.659 | n = 30 |
| Mother ID | 0.66 | 0.58 | 0.742 | n = 819 |

Supplementary Table 3: Bayesian GLMM estimates for August body weight in lambs. The sample size was n = 2286. The table shows the posterior mean and 95% credible intervals and information on how each variable was encoded and transformed prior to modelling.
